# RevGadgets: an R Package for visualizing Bayesian phylogenetic analyses from RevBayes

**DOI:** 10.1101/2021.05.10.443470

**Authors:** Carrie M. Tribble, William A. Freyman, Michael J. Landis, Jun Ying Lim, Joëlle Barido-Sottani, Bjørn Tore Kopperud, Sebastian Höhna, Michael R. May

## Abstract

1. Statistical phylogenetic methods are the foundation for a wide range of evolutionary and epidemiological studies. However, as these methods grow increasingly complex, users often encounter significant challenges with summarizing, visualizing, and communicating their key results.
2. We present `RevGadgets`, an `R` package for creating publication-quality figures from the results of a large variety of phylogenetic analyses performed in `RevBayes` (and other phylogenetic software packages).
3. We demonstrate how to use `RevGadgets` through a set of vignettes that cover the most common use cases that researchers will encounter.
4. `RevGadgets` is an open-source, extensible package that will continue to evolve in parallel with `RevBayes`, helping researchers to make sense of and communicate the results of a diverse array of analyses.

[Bayesian phylogenetics, data visualization, R, RevBayes]

## Introduction

Beyond being a graphical representation of the Tree of Life, phylogenetic trees provide a rigorous basis for a wide range of evolutionary and epidemiological inferences. Phylogenetic methods allow researchers to understand how molecular and morphological traits evolve (Nei, 1987; Yang, 2014; Felsenstein, 1985; Harvey and Pagel, 1991), how lineages disperse over geographic space (Ronquist and Sanmartín, 2011), and how lineages diversify over time (Morlon, 2014), among other evolutionary phenomena. Additionally, phylogenetic methods can be used to inform conservation decisions (Faith, 1992) and are powerful epidemiological tools (Volz et al., 2013; Baele et al., 2017).

Phylogenetic methods are increasingly based on explicit probabilistic models with parameters that describe underlying evolutionary processes. As datasets grow and evolutionary hypotheses become more nuanced, these models necessarily become more complex. `RevBayes` (Höhna et al., 2016) is a Bayesian phylogenetic inference program that was developed to accommodate this increasing complexity and allows users to explore a vast space of phylogenetic models. Models in `RevBayes` are specified as probabilistic graphical models (Höhna et al., 2014), which are graphical representations of the underlying dependencies among parameters (and their corresponding prior distributions), similar to individual Legos being used to build a complex city. Using this graphical modeling framework, users can design customized models and tailor analyses to their particular datasets and research questions. However, this flexibility comes at a cost: because of the nearly infinite variety of possible models (and model combinations) that users can explore in `RevBayes`, the results of these analyses are often challenging to summarize and visualize using standard software. This is a significant limitation for `RevBayes` users because, in addition to being the primary method for reporting results of phylogenetic analyses, graphical summaries are a valuable tool for making sense of scientific results (Tufte, 2001), and for diagnosing modeling and analytical problems (Kerman et al., 2008).

Historically, `RevBayes` users have had to process and plot their results using *ad hoc* scripts written for each analysis, which imposed a significant barrier to entry for users not familiar with the structure of `RevBayes` output or comfortable with developing their own graphical summaries. To address these challenges, we developed `RevGadgets. RevGadgets` is an `R` package (R Core Team, 2020) that adds to the diverse ecosystem of phylogenetic visualization tools—*e.g.*, `ape` (Paradis and Schliep, 2019), `Tracer` (Rambaut et al., 2018), `phytools` (Revell, 2012), `ggtree` (Yu et al., 2017), `FigTree` (Rambaut, 2014), `IcyTree` (Vaughan, 2017), among many others— but is specialized for output produced by `RevBayes. RevGadgets` serves as a bridge between `RevBayes` analyses and existing tools for phylogenetic data processing and plotting in `R`, especially the `ggtree` package suite, which includes the `ggtree, tidytree`, and `treeio` packages (Wang et al., 2020; Yu et al., 2017). `RevGadgets` provides tools for plotting summary trees (including summaries of parameters for each branch), ancestral-state estimates, and posterior distributions of parameters for a variety of models. Using the general framework of `ggplot2, the tidyverse`, and associated packages (Wickham, 2011; Wickham et al., 2019), plotting functions return plot objects with default, but customizable, aesthetics. Here, we present five vignettes demonstrating how to use `RevGadgets` to summarize results for a variety of phylogenetic analyses.

### Phylogenies

Phylogenies are central to all analyses in `RevBayes`, so accurate and information-rich visualizations of evolutionary trees are critical. In this case study, we demonstrate the tree-plotting functionality of `RevGadgets`, with methods to visualize phylogenies and their associated posterior probabilities, divergence-time estimates, and branch-specific parameter estimates.

`RevGadgets` provides paired functions for (1) reading in and processing data, and (2) summarizing and visualizing results. For phylogenies, the function `readTrees()` loads trees (either individual trees, or sets of trees) in either Newick or `NEXUS` (Maddison et al., 1997) formats, then processes associated branch or node annotations, and finally stores the tree(s) as `treedata` object(s) (as defined by `treeio`; Wang et al., 2020). Users can then visualize the `treedata` object using either `plotTree()` or `plotFBDTree()`, as we demonstrate below. Alternatively, users may choose to write custom plotting code using existing `ggtree` functions.

`RevGadgets` can plot both unrooted and rooted trees, and creates plots that are compatible with plotting options from `ggtree`. Additionally, `RevGadgets` provides extensive functionality for plotting trees with non-contemporaneous tips, such as those produced by total-evidence analyses under the fossilized birth-death [FBD] process (Heath et al., 2014; Zhang et al., 2016). The fossilized birth-death process (and the related serially-sampled birth-death process; Stadler, 2010) produces sampled ancestors (samples that are directly ancestral to another sampled taxon and thus are not represented as tips in the tree), and the ages of the samples are often subject to uncertainty (*e.g.*, because of imperfect knowledge about the age of the strata from which the samples were collected). As a consequence, conventional tree plotting tools are unsuitable for plotting FBD trees. We demonstrate how to use `RevGadgets` to plot the results of an FBD analyses of living and extinct bears (Figure 1; data from Abella et al., 2012 and Heath et al., 2014). We include age bars colored by the posterior probability of the corresponding node, a geological time scale and labeled epochs from the package `deeptime` (Gearty, 2021), and fossils estimated to be direct ancestors of other samples (*i.e.*, sampled ancestors).

**Figure 1:**
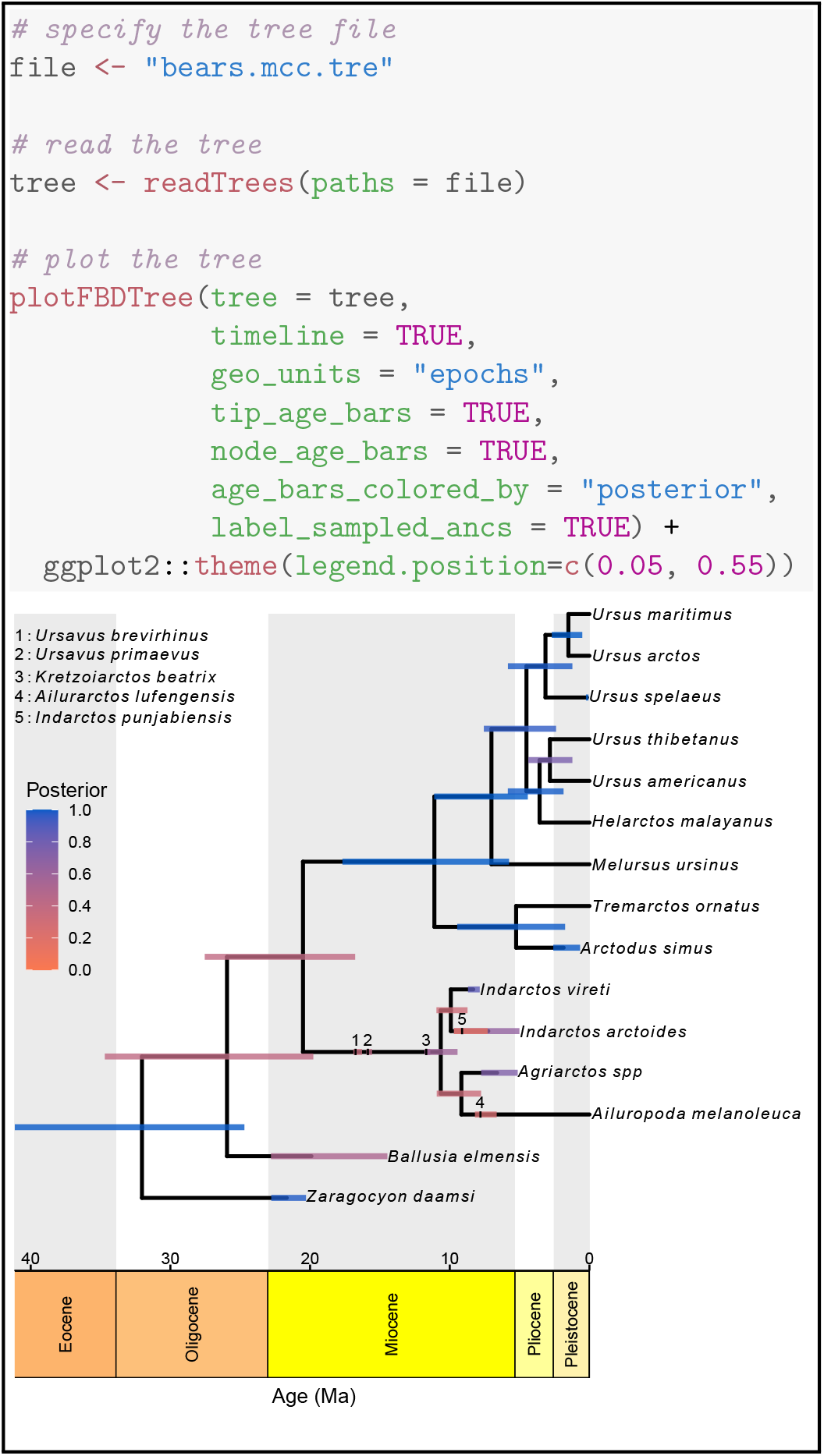
Plotting a time-calibrated phylogeny of extinct and extant taxa. Top) `RevGadgets` code for reading in and plotting a time-calibrated phylogeny of extant and extinct bears. We use the theme function from `ggplot2` to add the posterior-probability legend. Bottom) The maximum sampled-ancestor clade-credibility (MSACC) tree for the bears. Sampled ancestors are indicated by numbers along the branches (legend, top left). Bars represent the 95% credible interval of the age of the node, tip or sampled ancestor in millions of years (geological timescale, x-axis); the color of the bar corresponds to the posterior probability (legend, middle left) of that a clade exists, the posterior probability that a fossil is a sampled ancestor, or the posterior probability that a tip is not a sampled ancestor. (Data from Abella et al., 2012; Heath et al., 2014.)

In addition to visualizing trees themselves, `RevGadgets` allows researchers to visualize branchspecific parameters, for example rates of evolution or diversification for each branch in the phylogeny. In Figure 2, we demonstrate how to use `plotTree()` to visualize the estimated optimal body size as it varies across the cetacean phylogeny under a relaxed Ornstein-Uhlenbeck process (Butler and King, 2004; Uyeda and Harmon, 2014; data from Steeman et al., 2009; Slater et al., 2010). Under this model, a quantitative character evolves towards an adaptive optimum that changes along the branches of the tree, and thus the optimum associated with each branch is a focal inference.

**Figure 2:**
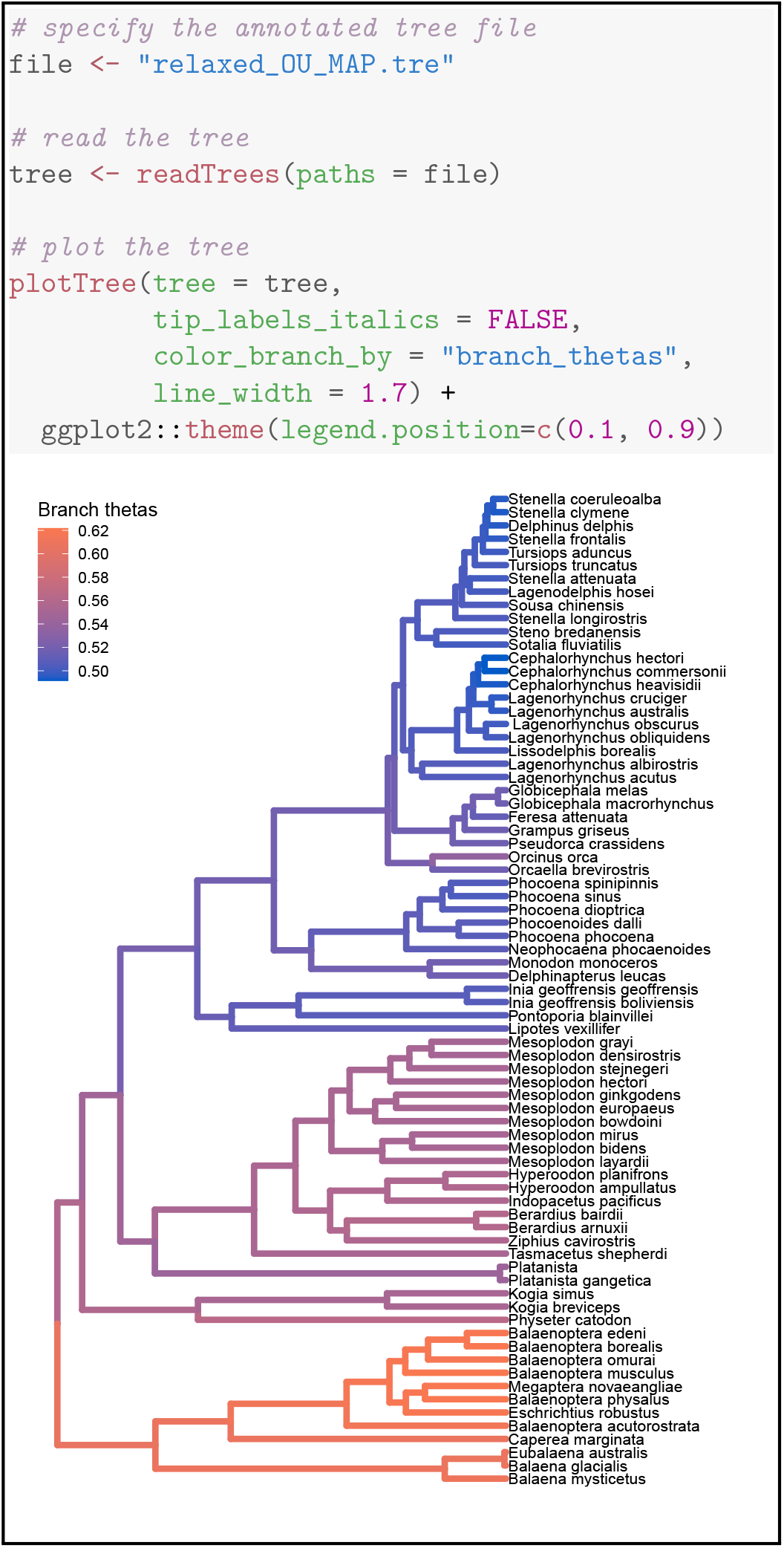
Plotting branch-specific parameter values across a phylogeny. Top) `RevGadgets` code for reading in and plotting the cetacean phylogeny that has been annotated with branch-specific adaptive optima (*θ*) inferred under a relaxed Ornstein-Uhlenbeck model. Bottom) The cetacean phylogeny with branches colored according to the posterior-mean estimate of the inferred branch-specific optimum body size, *θ* (legend, tip left). (Phylogeny from Steeman et al., 2009; body size data in units of natural log-transformed meters from Slater et al., 2010.)

The `plotTree()` function can also visualize unrooted or circular phylogenies, and users may add text annotations to denote posterior probabilities or other quantities. Users can apply `ggtree` functions to modify the `RevGadgets` plot, *e.g.*, to highlight certain clades with `geom hilight()` or to add phylopics (http://phylopic.org/) using `geom phylopic()`. Together, these functions provide user-friendly and customizable tree-plotting functionality for a variety of core research questions in evolutionary biology.

### Posterior Estimates of Numerical Parameters

`RevGadgets` provides several tools to visualize posterior distributions of numerical parameters. The output produced by most `RevBayes` analyses is a (typically tab-delimited) text file where rows correspond to samples from sequential iterations of an MCMC analysis, and columns correspond to parameters in the model. Most information of interest to researchers—*e.g.*, most probable parameter values (maximum *a posteriori*, or MAP, estimates), 95% credible intervals (CIs), or full posterior distributions—requires processing this raw MCMC output. Here, we demonstrate methods for processing and visualizing MCMC output for both quantitative and qualitative parameters.

We illustrate the core functions for reading, summarizing and visualizing posterior distributions of specific parameters with an example analysis of chromosome number evolution (Figure 3; data from Freyman and Höhna, 2018). We use `readTrace()` to read in parameters sampled during one or more MCMC analyses. We then use `summarizeTrace()` to calculate the posterior mean and 95% credible interval for the focal parameters. Finally, we plot the marginal posterior distributions of the focal parameters using `plotTrace()`.

**Figure 3:**
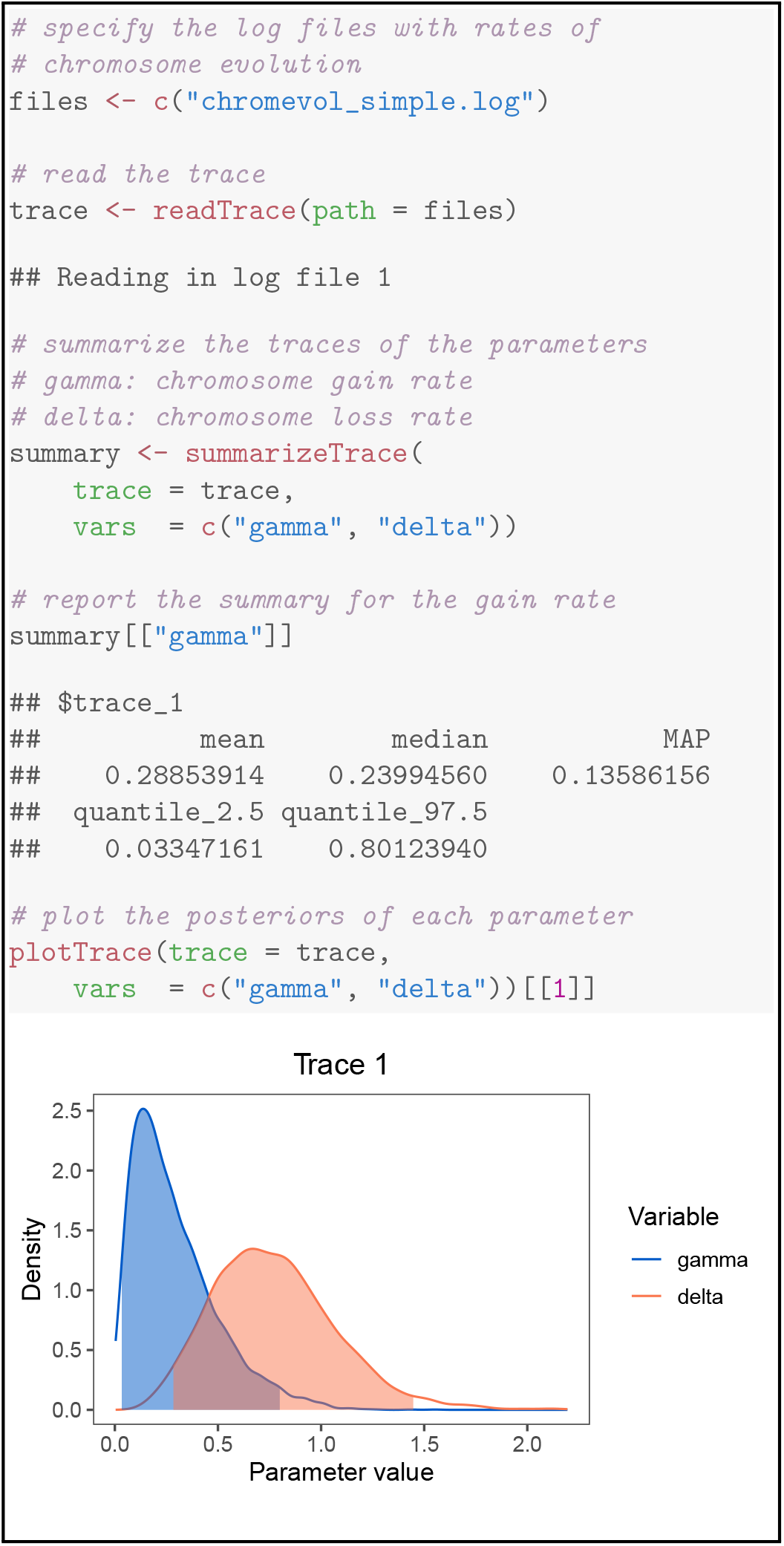
Plotting posterior distributions of numerical parameter values. Top) `RevGadgets` code for reading in and plotting the posterior distributions of rates of chromosome evolution in *Aristolochia*. Bottom) Marginal posterior distributions of the two rate parameters. Shaded regions represent the 95% credible interval of each posterior distribution. (Data from Freyman and Höhna, 2018.)

Plots of the posterior distributions of parameter values are key to a thorough understanding of the results of any Bayesian analysis. These tools encourage users to explore their results thoroughly rather than relying on single summary statistics. These summaries and plots may also be useful as tools for science communication and education on statistical phylogenetics, as they can easily be used to demonstrate differences in parameter estimates that result from changes to basic phylogenetic models. Additionally, the output of `readTrace()` may be passed to `R` packages specializing in MCMC diagnosis, *e.g.*, `convenience` (Fabreti and Höhna, 2021) or `coda` (Plummer et al., 2006). These functions are compatible with any delimited text file of MCMC samples, and can be used with the output of most Bayesian phylogenetic programs.

### Ancestral-State Estimates

In addition to making inferences about the underlying process of evolution, researchers may be interested in studying how particular characters evolved across the branches of the phylogeny. Ancestral-state estimation is a method for inferring that history.

`RevGadgets` offers two different types of summaries for ancestral-state estimates: 1) maximum *a posterior* (MAP) estimates, *i.e.*, the states with the highest posterior probability at each node, and; 2) pie charts that represent each state in proportion to its probability at each node. Ancestral-state estimates may be represented as text annotations rather than colored symbols. Additionally, `RevGadgets` can summarize and visualize ancestral-state estimates at internal nodes and at the “shoulders”, *i.e.*, at the beginning of each branch. Plotting the states at internal nodes is appropriate for standard evolutionary models of anagenetic (within-lineage) change. However, models of evolution that include a cladogenetic component (*e.g.*, models of biogeographic or chromosome-number evolution; Ree and Smith, 2008; Goldberg and Igić, 2012; Freyman and Höhna, 2018) also allow states to change at speciation events. In this case, researchers may also want to plot the shoulder states, which represent the ancestral-state estimates for each daughter lineage immediately following the speciation event.

We demonstrate how to plot ancestral-state estimates of placenta type across the mammal phylogeny under an asymmetric model of character evolution (Figure 4; data from Elliot and Crespi, 2006). First, we use `processAncStates()` to read in and parse the phylogeny and ancestral-state estimates inferred using `RevBayes`. Second, we use `plotAncStatesMAP()` to color each node symbol according to the state with the highest posterior probability, and make the area of the symbol proportional to that state’s posterior probability. Because of the size of the phylogeny, we choose to plot the estimates on a circular tree by changing the tree layout parameter.

**Figure 4:**
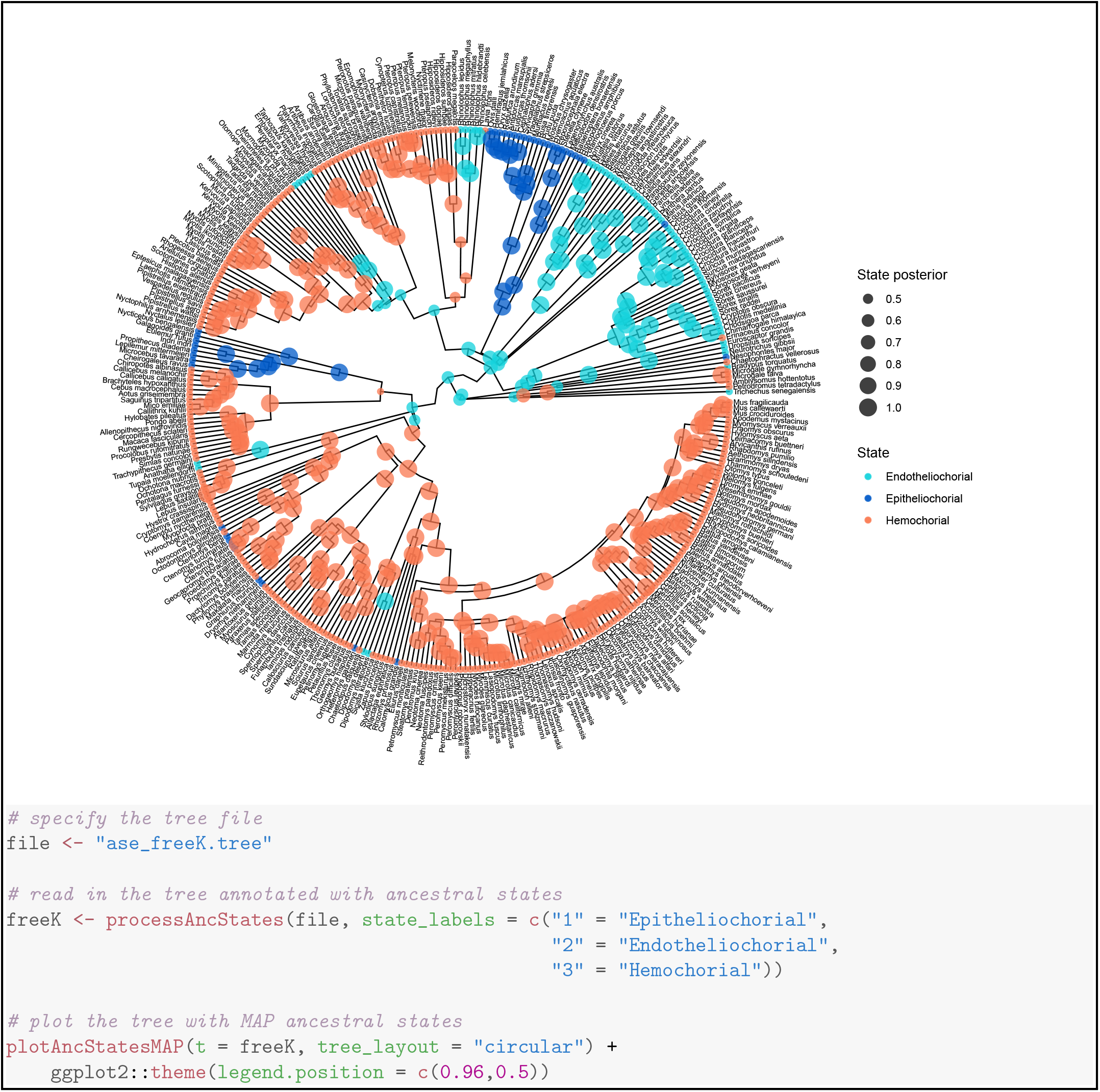
Plotting maximum *a posterior* (MAP) estimates of ancestral states on a circular phylogeny. Top) MAP estimates of ancestral placental states across the phylogeny of mammals. Each node is colored by the MAP state (legend, bottom right); the size of each symbol is proportional to the posterior probability of the map state (legend, top right). Bottom) `RevGadgets` code for reading in and plotting the MAP estimates for ancestral placental states across the mammals phylogeny. (Data from Elliot and Crespi, 2006.)

Next, we demonstrate plotting estimates of ancestral ranges of the Hawaiian silversword alliance that were generated by a Dispersal-Extinction-Cladogenesis (DEC) model (Figure 5; data from Landis et al., 2018). Since the DEC model features a cladogenetic component, we include shoulder-state estimates. Because of the large number of states in this analysis (15 possible ranges and one “other” category), more pre-processing is necessary. As before, we pass the appropriate state names to `processAncStates()`; however, in this case we plot pie charts representing the probability of each state using `plotAncStatesPie()`, and plot states at shoulders using `cladogenetic = TRUE`.

**Figure 5:**
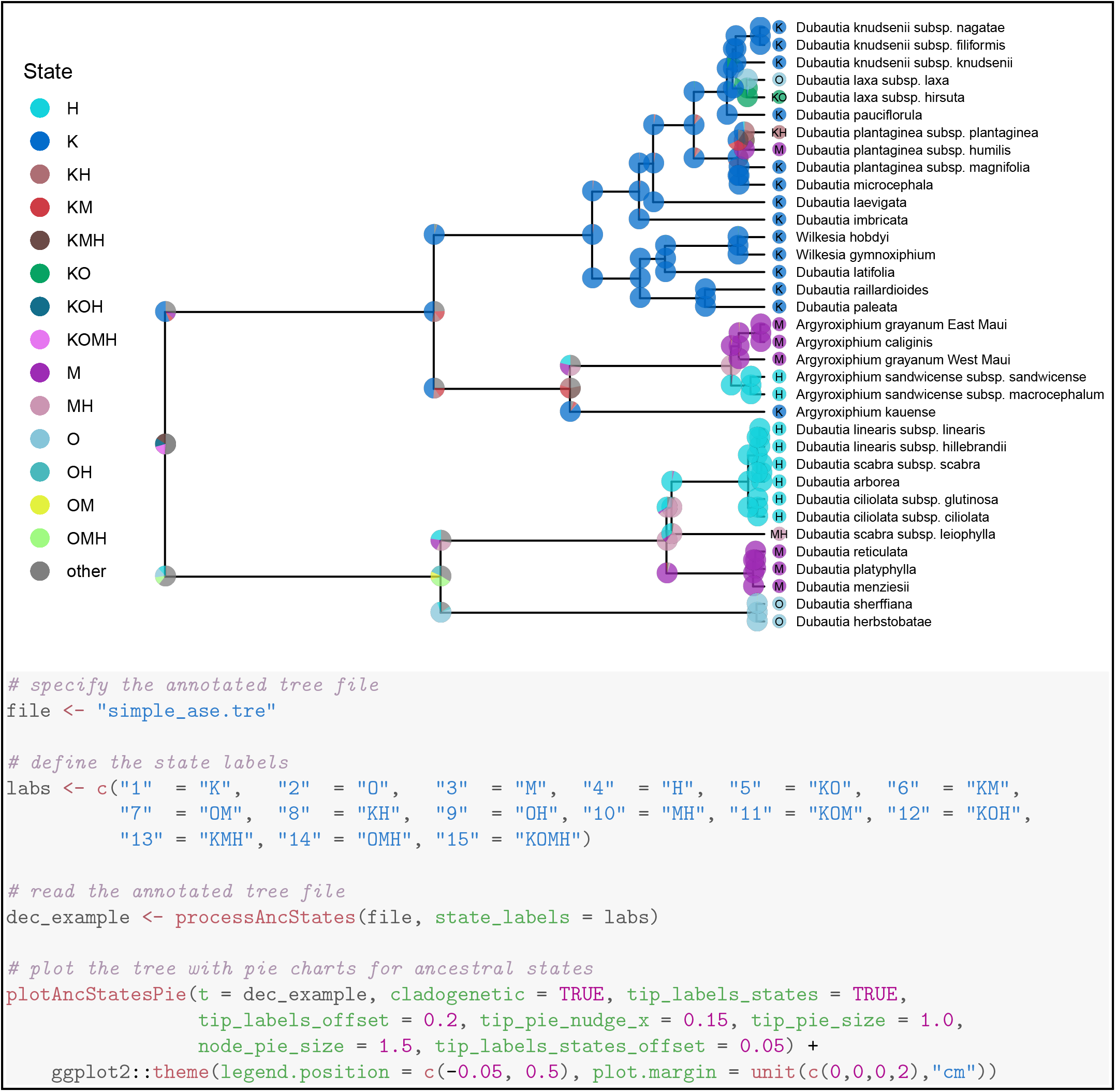
Plotting posterior distributions of ancestral states under a cladogenetic model. Top) The posterior estimates ancestral biogeographic states of the Hawaiian silverswords estimated under a DEC model. The size of each pie slice is proportional to the posterior probability of a given state (legend, top left) for a particular lineage. Pies at nodes represent the state of the ancestral lineage immediately before speciation; pies at “shoulders” represent the states of each daughter lineage immediately following the speciation event. Bottom) `RevGadgets` code for reading in and plotting the posterior estimates for ancestral geographic range across the phylogeny of Hawaiian silverswords. (Data from Landis et al., 2018.)

Beyond the above examples, these versatile plotting tools can visualize any discrete ancestral-state estimates reconstructed by `RevBayes`, including the results of chromosome count estimations (Freyman and Höhna, 2018) and discrete state-dependent speciation and extinction (SSE) models (Freyman and Höhna, 2019; Zenil-Ferguson et al., 2019).

### Diversification Rates

The processes of speciation and extinction (*i.e.*, lineage diversification) is of great interest to evolutionary biologists (Morlon, 2014). Rates of speciation and extinction may be modeled as constant over time and among branches (as in a constant-rate birth-death process; Kendall et al., 1948; Nee et al., 1994), or allowed to vary over time (Stadler, 2011; May et al., 2016), across branches of a phylogeny (Rabosky, 2014; Höhna et al., 2019), or based on the character states of the evolving lineages (Maddison et al., 2007; Freyman and Höhna, 2019). For example, rates that vary across branches of the phylogeny can be visualized using `plotTree()` to color the branches by their inferred rate. State-dependent diversification models provide estimates of the speciation and extinction rates associated with each character state, and may also be used to estimate ancestral states. `plotTrace()` or specific processing and plotting functions for diversification rates—`processSSE(), plotMuSSE`, and `plotHiSSE`—may be used to visualize the estimated rates. `plotAncStatesMAP()` or `plotAncStatesPie()` may be used to visualize the ancestral-state estimates.

We demonstrate how to plot the results of a time-varying model—the episodic birth-death process (Stadler, 2011; Höhna, 2015)—applied to primate phylogeny (Figure 6; Springer et al., 2012). The episodic birth-death analysis in `RevBayes` produces separate trace files each type of rate. We read these output files using `processDivRates()` and plot the resulting parameter estimates over time using `plotDivRates()`.

**Figure 6:**
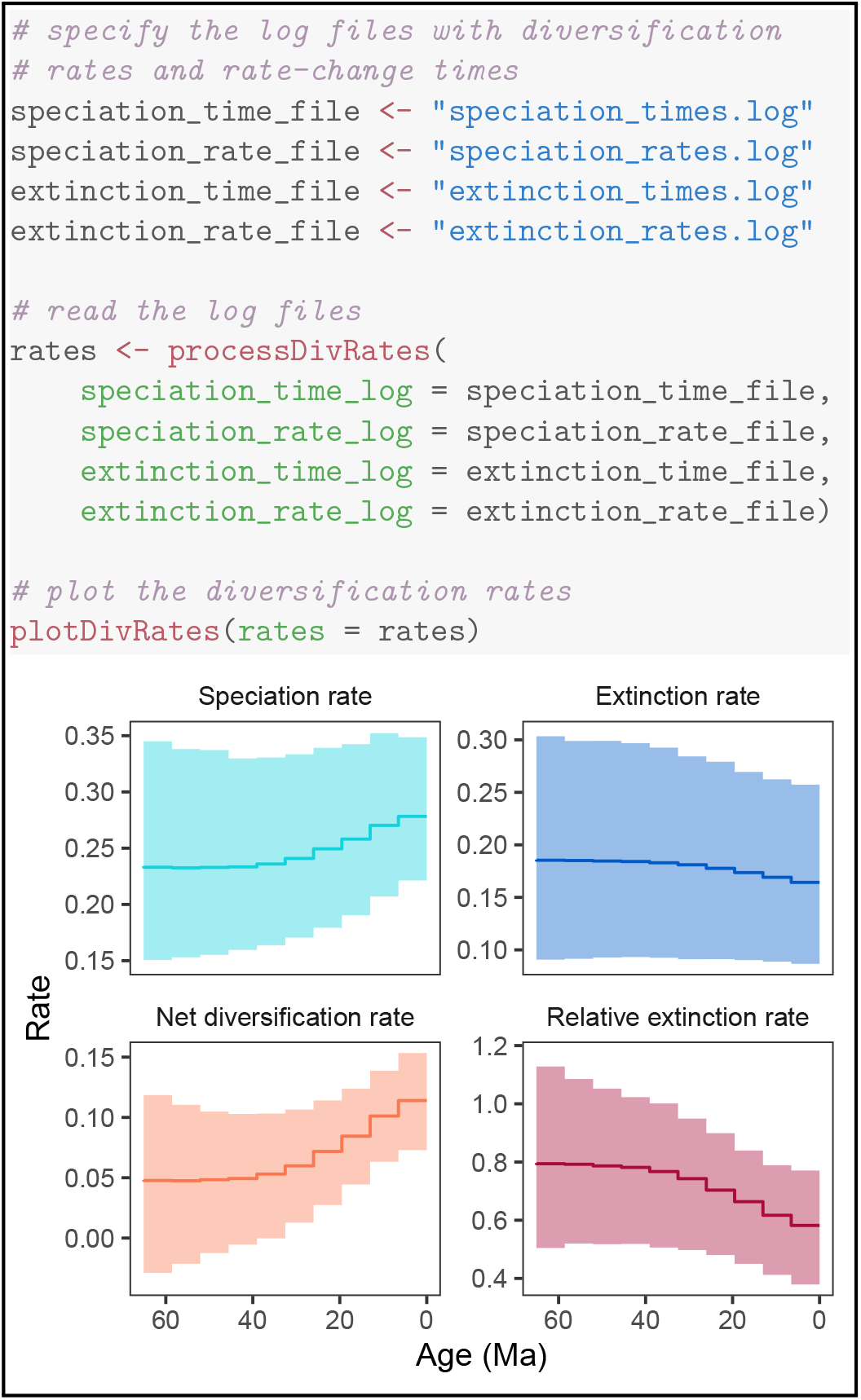
Plotting posterior distributions of diversification rates over time. Top) `RevGadgets` code for reading in and plotting the posterior estimates of diversification rates over time inferred from the primate phylogeny. Bottom) Posterior distributions of speciation and extinction rates over time, as well as the net diversification rate (speciation minus extinction) and the relative extinction rate (extinction divided by speciation). Dark lines correspond to the posterior-mean estimate of each parameter for each time interval, and shaded regions correspond to the 95% credible interval. (Data from Springer et al., 2012.)

Together with the aforementioned functions for plotting diversification parameter estimates, `plotDivRates()` allows users to visualize the outputs of nearly all diversification analyses available in `RevBayes`. Stochastic character mapping of diversification estimates, in which the timing and location of diversification rate shifts are painted along the branches of the tree, will be added in the future (Freyman and Höhna, 2019; Höhna et al., 2019).

### Model Adequacy

In addition to visualizing the results of phylogenetic inferences with a specific model, `RevGadgets` provides tools for exploring the adequacy of the model (*i.e.*, whether the model provides an adequate description of the data-generating process; Bollback, 2002; Gelman et al., 2013; Brown, 2014; Höhna et al., 2018). Posterior-predictive analysis tests whether a fitted model simulates (predicts) data that are similar to the observed data. This process is distinct from model testing, in which one model is chosen from a set of possible models, as the best model of the set may still provide an inadequate description of the underlying process.

First, users analyze their data with the model of interest and then use the inferred posterior distribution to simulate a number of new data sets. The user then selects test statistics that describe important features of the data (*e.g.*, the number of invariant sites in a nucleotide alignment) and calculates these statistics for both the observed data and the simulated data. If the statistic from the empirical data is reasonably included within the distribution of statistics from simulated datasets (posterior-predictive *p*-value > 0.05), the model is considered an adequate description of the process that produced the tested data feature.

Here, we demonstrate the workflow for a posterior-predictive analysis to test model adequacy of the Jukes-Cantor model for nucleotide sequence evolution (Jukes et al., 1969) in a single gene across a sample of 23 primates (Figure 7; data from Springer et al., 2012). First, we perform an analysis in `RevBayes` under a Jukes-Cantor model of nucleotide sequence data. Second, we use `RevBayes` to simulate datasets under the posterior distributions estimated in the first step. Third, we use `RevBayes` to calculate statistics from the simulated and empirical datasets. These statistics should describe aspects of the data that we hope capture a meaningful component of model performance. Finally, we use `RevGadgets` to plot those statistics and compute posterior-predictive *p*-values.

**Figure 7:**
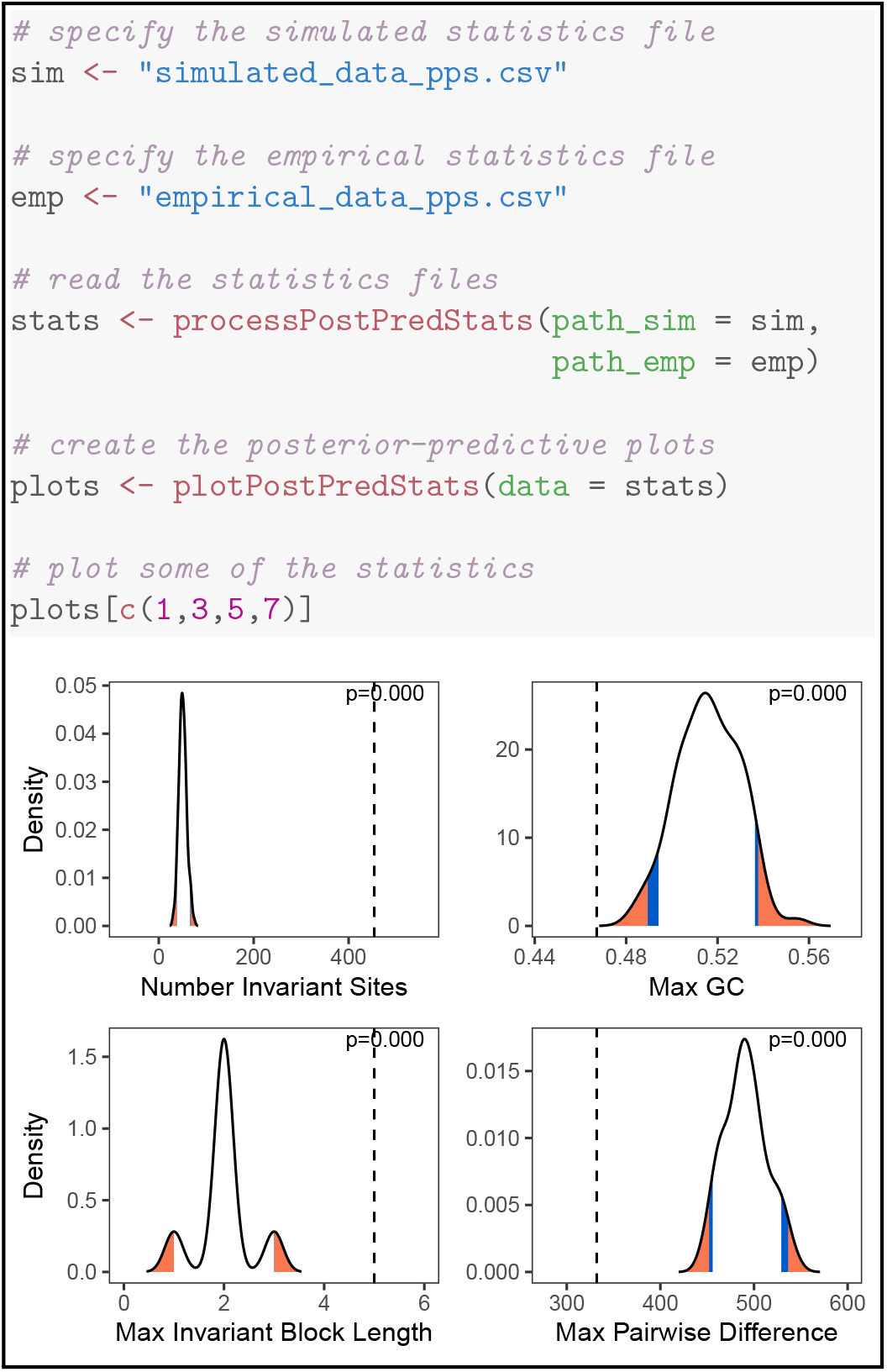
Plotting simulated posterior-predictive distributions to assess model adequacy. Top) `RevGadgets` code for reading in and plotting the distributions of summary statistics generated using posterior-predictive simulation posterior. Bottom) Posterior-predictive distributions (black curves) of four statistics simulated under the Jukes-Cantor model fit to primate *cytb*, compared to the same statistics computed on the observed data (dashed vertical lines). The posterior-predictive *p*-value (upper right of each panel) is the fraction of simulated statistics that are as or more extreme than the observed statistic. If the observed statistic falls in or beyond the orange region, we deem the model as inadequate at the 5% significance level; if the observed statistic falls in the blue region, the model is marginally adequate at the 10% significance level. In this case, the Jukes-Cantor model provides an inadequate description of the true generating process according to every summary statistic. (Data from Springer et al., 2012.)

Despite being computationally inexpensive compared to Bayesian model comparison methods (*i.e.*, Bayes factor calculation), posterior-predictive approaches remain relatively uncommon in empirical phylogenetic studies. As genome-scale datasets and increasingly complex statistical methods become more accessible to researchers, posterior-predictive simulation will be critical to testing how well our models describe the underlying generative processes. This component of `RevGadgets` functionality and the associated clear workflows for performing and interpreting posterior-predictive tests will hopefully increase the adoption of this important tool.

## Conclusions

`RevBayes` is a flexible platform for performing Bayesian phylogenetic evolutionary inferences. Because of the almost endless possibilities for building unique combinations of models in `RevBayes`, these analyses are often challenging to visualize using standard plotting software. We have developed an `R` package, `RevGadgets`, to produce publication-quality visualizations of phylogenetic analyses performed in `RevBayes`. The case studies described above illustrate some of the core functionality available in `RevGadgets` and demonstrate how to produce plots of the most commonly-performed `RevBayes` analyses. `RevBayes` is open source software that is actively maintained and developed. Likewise, `RevGadgets` is also open source and will continue to provide new plotting tools to meet new visualization challenges as they arise. `RevGadgets` and any future updates will be available on CRAN (https://cran.r-project.org/web/packages/RevGadgets/index.html) and on GitHub at https://github.com/cmt2/RevGadgets. Additionally, we provide thorough documentation for all functionality in the package and maintain numerous tutorials demonstrating how to use `RevGadgets` on the `RevBayes` website at https://revbayes.github.io/tutorials/. Together, the modular modeling tools from `RevBayes` and the visualization gadgets in `RevGadgets` will help researchers make sense of and communicate the results of a diverse array of sophisticated phylogenetic analyses.

## Authors Contributions

CMT and MRM designed the R package. All authors contributed code and examples. CMT and MRM drafted the manuscript. All authors revised and approved the final version of the manuscript.

## Dependencies

`RevGadgets` depends on many `R` packages, in particular: `ape` (Paradis and Schliep, 2019); `phangorn` (Schliep, 2011); `phytools` (Revell, 2012); `ggplot2` (Wickham, 2011); `ggtree` (Yu et al., 2017); `treeio` (Wang et al., 2020); `deeptime` (Gearty, 2021); `dplyr` (Wickham et al., 2021); `treeplyr` (Uyeda and Harmon, 2020); `tidytree` (Yu, 2021b); `reshape` (Wickham, 2007); `ggthemes` (Arnold, 2021); `tidyr` (Wickham, 2021); `tibble` (Müller and Wickham, 2021); `gginnards` (Aphalo, 2021a); `ggimage` (Yu, 2020); `ggplotify` (Yu, 2021a); `png` (Urbanek, 2013); and `ggpp` (Aphalo, 2021b).

## Acknowledgements

We would like to acknowledge Carl J. Rothfels, Benjamin K. Blackman, David D. Ackerly, and Chelsea D. Specht for feedback on initial stages of the manuscript. Ixchel González Ramírez, Jenna T. B. Ekwealor, Isaac Lichter Marck, and members of the Rothfels Lab at UC Berkeley provided valuable feedback on usability and legibility of figures and code. Klaus Schliep and an anonymous reviewer provided important feedback on the package structure and stability. Andrew Magee, Kengo Nagashima, Klaus Schliep, and Josef Uyeda contributed code.

This research was supported by the Deutsche Forschungsgemeinschaft (DFG) Emmy Noether-Program HO 6201/1-1 awarded to SH.

## Data Availability

`RevGadgets` is hosted on CRAN (https://cran.r-project.org/web/packages/RevGadgets/index.html) and all example datasets are freely available on the RevBayes website (https://revbayes.github.io/tutorials/intro/revgadgets).

## Notes

https://github.com/cmt2/RevGadgets

### Competing Interest Statement

The authors have declared no competing interest.

### Summary of Updates

RevGadgets has now been made available on CRAN

